# Identification of key pathways and metabolic fingerprints of longevity in *C. elegans*

**DOI:** 10.1101/222554

**Authors:** Arwen W. Gao, Reuben L. Smith, Michel van Weeghel, Rashmi Kamble, Riekelt H. Houtkooper

## Abstract

Impaired insulin/IGF-1 signaling (IIS) pathway and caloric restriction (CR) are two well-established interventions to prolong lifespan in worm *C. elegans*. However, a cross comparison of these longevity pathways using a multi-omics integration approach is lacking. In this study, we aimed to identify key pathways and metabolite fingerprints of longevity that are shared between IIS and CR worm models using a multi-omics integration approach. We generated transcriptomics and metabolomics data from two long-lived mutant worm strains, i.e. *daf-2* (impaired IIS pathway) and *eat-2* (CR model) and compared them with the N2 strain. Transcriptional profiling identified shared longevity signatures between the two strains, such as an upregulation of lipid storage and defense responses and downregulation of macromolecule synthesis and developmental processes. The shared longevity signatures revealed by metabolomics profiling included an increase in the levels of glycerol-3P, ademine, xanthine, and AMP, and a decrease in the levels of the amino acid pool, the C18:0 and C17:1 fatty acids. After we integrated transcriptomics and metabolomics data based on the annotations in KEGG, our results highlighted a downregulation of pyrimidine metabolism and upregulation of purine metabolism as a commonality between the two longevity mechanisms. Overall, our findings point towards the existence of shared metabolic pathways that are likely important for lifespan extension and provide novel insights into potential regulators and metabolic fingerprints for longevity.

## Introduction

Average life expectancy has dramatically increased over the past few decades, resulting from improvements in healthcare systems in most economically developed countries^1^. Aging and life expectancy are effected by genetic predisposition and non-genetic and environmental risk factors, making the process of aging different between individuals^2^. Although the genetic and environmental causes of aging are not fully established, several processes have been identified as hallmarks of aging. Many of these hallmarks involve metabolism, such as deregulated nutrient sensing and mitochondrial dysfunction^2^. A prototypical example of how environmental factors are involved in the metabolic network of longevity is caloric stress. One example of this is caloric restriction (CR), a condition of 20-50% reduced caloric intake, that activates several metabolic pathways and thereby extends lifespan across species^3^. Besides environmental factors, genetics also play an important role in the metabolic control of aging. So far, several metabolic genes have been identified that influence longevity^3^. For instance, mammalian longevity is influenced by metabolic changes as a result of genetically determined insulin sensitivity and glucose handling^4^. Taken together, it is evident that both the environment and genetics play an important role in controlling metabolism and aging. However, it remains to be elucidated how both components interact to regulate metabolism and aging.

*C. elegans* is one of the most popular model organisms for aging studies due to its relatively short lifespan, fully annotated genome and its ease of manipulation^5^. The first aging gene to be discovered was *daf-2*, which encodes a homolog of mammalian insulin/insulin-like growth factor (IGF), and its regulatory mechanisms involved in longevity are highly conserved across a wide range of species, including yeast, flies, mice and human beings^6^. Lowered DAF-2 signaling leads to translocation of DAF-16, a fork-head transcription factor, that enters the nucleus and activates the expression of numerous genes involved in longevity, lipid metabolism, stress response and innate immunity^7^. As a result, *daf-2* mutant animals can live twice as long as control animals^8^. In addition, *C. elegans* also serves as a tractable model to study molecular mechanisms underlying CR. Distinct fashions of CR exist, including genetic (i.e. *eat-2* mutants) and dietary interventions (e.g. diluting bacterial food)^9,10^. *eat-2* mutant animals have a dysfunctional pharynx, which slows down food intake, and results in a lifespan increase of approximately 50%^5^. Previous studies on *daf-2* and *eat-2* mutants have outlined the essential biological processes involved in the regulatory networks of aging, and have opened up novel avenues for treatments and biomarkers for aging-related diseases^7^. In fact, longevity mediated by CR and impaired insulin/IGF-1-like signaling (IIS) seems to be partially overlapping in terms of the downstream regulatory processes to mediate lifespan extension, including activation of autophagy, and inhibition of the target of rapamycin pathway (TOR)^7,11-13^. Although the IIS pathway, especially the activation of DAF-16, is associated with CR-induced longevity, it is dispensable for the longevity phenotype observed in *eat-2* mutants^7,9^.

In recent years, studies using high-throughput “omics” approaches have gained commendation as they have provided comprehensive information on the genomic, metabolomic, and proteomic changes that occur during the aging process^14^. Among all the omics analyses, metabolomics or profiling of metabolites, represents possibly the most diverse and complex level of biological regulatory processes, and is intimately linked to the phenotype. For instance, metabolite profiles facilitate the generation of phenotypic data of complex metabolic alterations that incorporate messages from multiple levels of systemic regulation, including the genome, the transcriptome, the proteome, the environment and their interactions^15^. Technological advancements in metabolomics studies have enabled numerous discoveries of perturbations in physiological networks and have further expanded our knowledge of longevity mechanisms underlying specific biological functions in different organisms, including worms, mice and humans^16-18^.

Although distinct age-related metabolic signatures have been reported in *C. elegans*, the relationship between gene transcription and metabolite levels remains to be fully elucidated. In this study, we aim to identify key metabolic signatures of longevity. We directly compared two long-lived mutant strains, namely *daf-2(e1370)* and *eat-2(ad465)* (from here on called *daf-2* and *eat-2* mutants) by whole-genome microarray and mass spectrometry-based platforms^18^. Next, we superimposed these two platforms in a cross-omics approach to visualize the actual changes at two regulatory levels, and identified shared and opposed modes of metabolic regulation of longevity.

## Results

### Gene expression analysis reveals the major signaling pathways important for longevity

To comprehensively quantify changes in global gene expression, we performed microarray analysis for the long-lived mutants *daf-2* and *eat-2* and used N2 worms as the control group. Differentially expressed genes (DEGs) in the long-lived mutants compared to N2, were analyzed using gene ontology (GO) and Kyoto Encyclopedia of Genes and Genomes (KEGG) pathway enrichment. In *daf-2* mutants, 3544 genes were significantly upregulated and 2336 genes were downregulated compared to those in N2 worms (Fig. 1a). In *eat-2* mutants, we found fewer genes to be significantly altered compared to *daf-2* mutants, 2352 and 983 genes were up- and down-regulated, respectively, compared to those in N2 worms (Fig. 1b).

**Figure 1.**
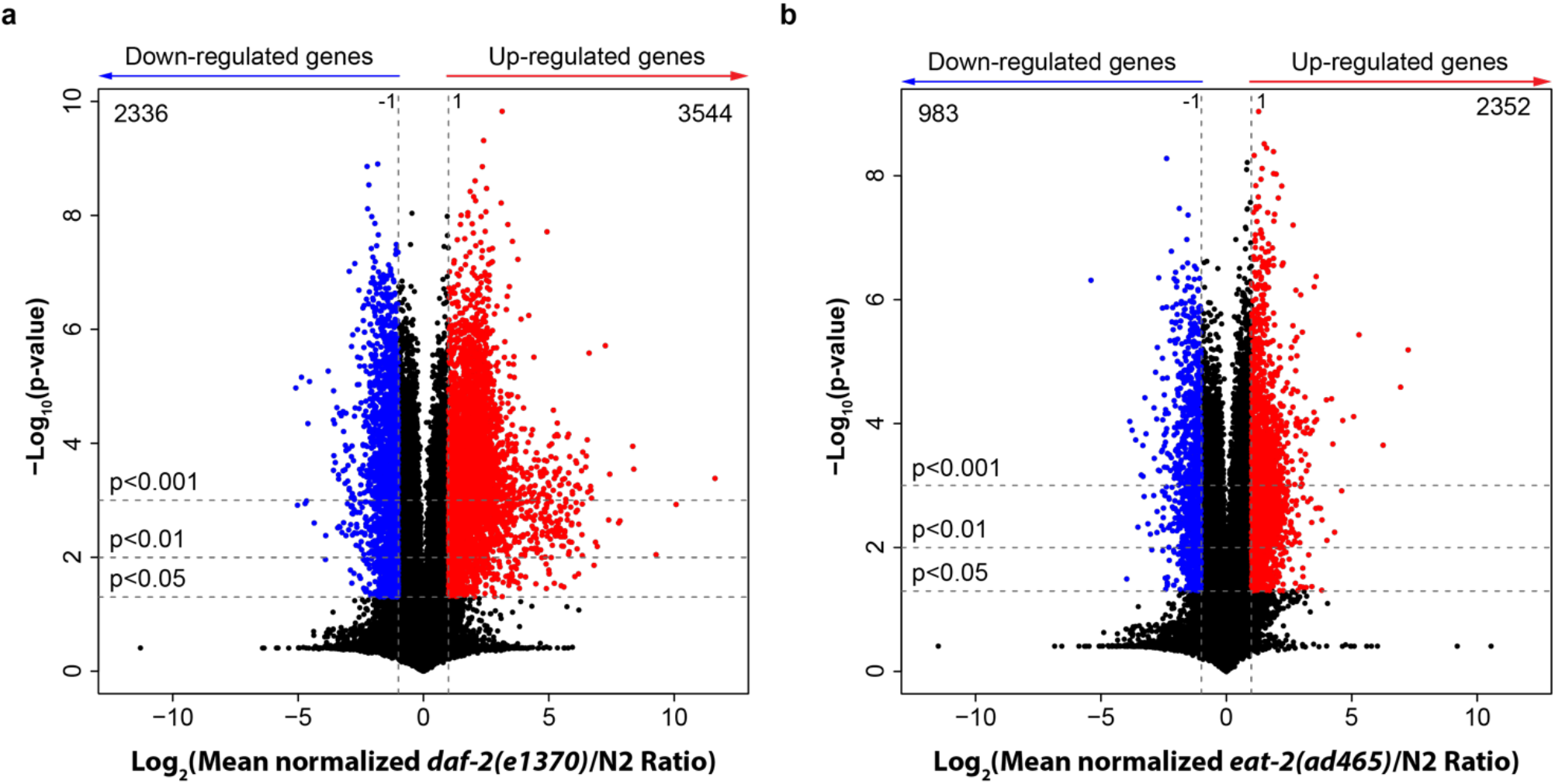
Transcriptional alterations in long-lived *daf-2* and *eat-2* mutants compared to N2 worms. (**a**) Volcano plot of log_2_-transformed fold change of gene expression in *daf-2(e1370)* over N2 worms. (**b**) Volcano plot of log_2_-transformed fold change of gene expression in *eat-2(ad465)* compared to those in N2 worms. p-value was calculated using student’s t-test. The vertical dashed lines represent the log_2_ fold change= −1 or 1, horizontal dashed lines denote p-value cutoffs of 0.05 (*), 0.01 (**), and 0.001 (***). The number of genes that are significantly up- (red) or down- (blue) regulated (p-value<0.05) in each panel is also mentioned.

To find shared pathways that are important for regulation of longevity, we focused on the overlapping genes between *daf-2* and *eat-2* mutants that have altered expression levels compared to N2 worms using the Database for Annotation, Visualization and Integrated Discovery (DAVID)^19^ and ReviGo^20^ (Fig. 2). GO term enrichment analysis for biological processes was performed on 1722 genes that were upregulated and 617 genes that were downregulated in both long-lived mutants compared to N2 worms. Enriched GO terms for upregulated genes included phosphorus metabolic process, lipid localization, regulation of anatomical structure, morphogenesis, cell surface receptor signaling pathways, response to stress, regulation of cell proliferation, male meiosis, spermatogenesis and flagellated sperm motility (Fig. 2a). For the 617 genes that were downregulated in both long-lived mutants, we observed an enrichment of GO terms for aromatic compound biosynthesis, heterocycle biosynthesis, RNA metabolism, regulation of RNA stability, response to abiotic stimulus, positive regulation of nucleobase-containing compound metabolism, regulation of nitrogen compound metabolism, mesoderm development and regulation of cell differentiation (Fig. 2b). Similar results were obtained when analyzing the upregulated genes for KEGG pathway analysis using DAVID (Table 1). We observed a significant enrichment in intermediary metabolism processes including fatty acid degradation, metabolic pathways, biosynthesis of antibiotics, metabolism of xenobiotics by cytochrome P450, drug metabolism, amino acid synthesis, lipid metabolism, tyrosine metabolism, glycine, serine and threonine metabolism, aromatic compound degradation, branched-chain amino acid synthesis, and cysteine and methionine metabolism. Downregulated genes mapped to the TGF-beta signaling pathway, Wnt signaling pathway, protein processing in the endoplasmic reticulum and ubiquitin mediated proteolysis pathways (Table 1). Taken together, we observed an enrichment of cellular metabolic pathways and energy metabolism pathways in both long-lived mutants. Furthermore, pathways that regulate the biosynthesis of macromolecules, such as aromatic and nitrogen compounds, were attenuated in both long-lived mutants, suggesting an overall low energy-consuming state.

**Figure 2.**
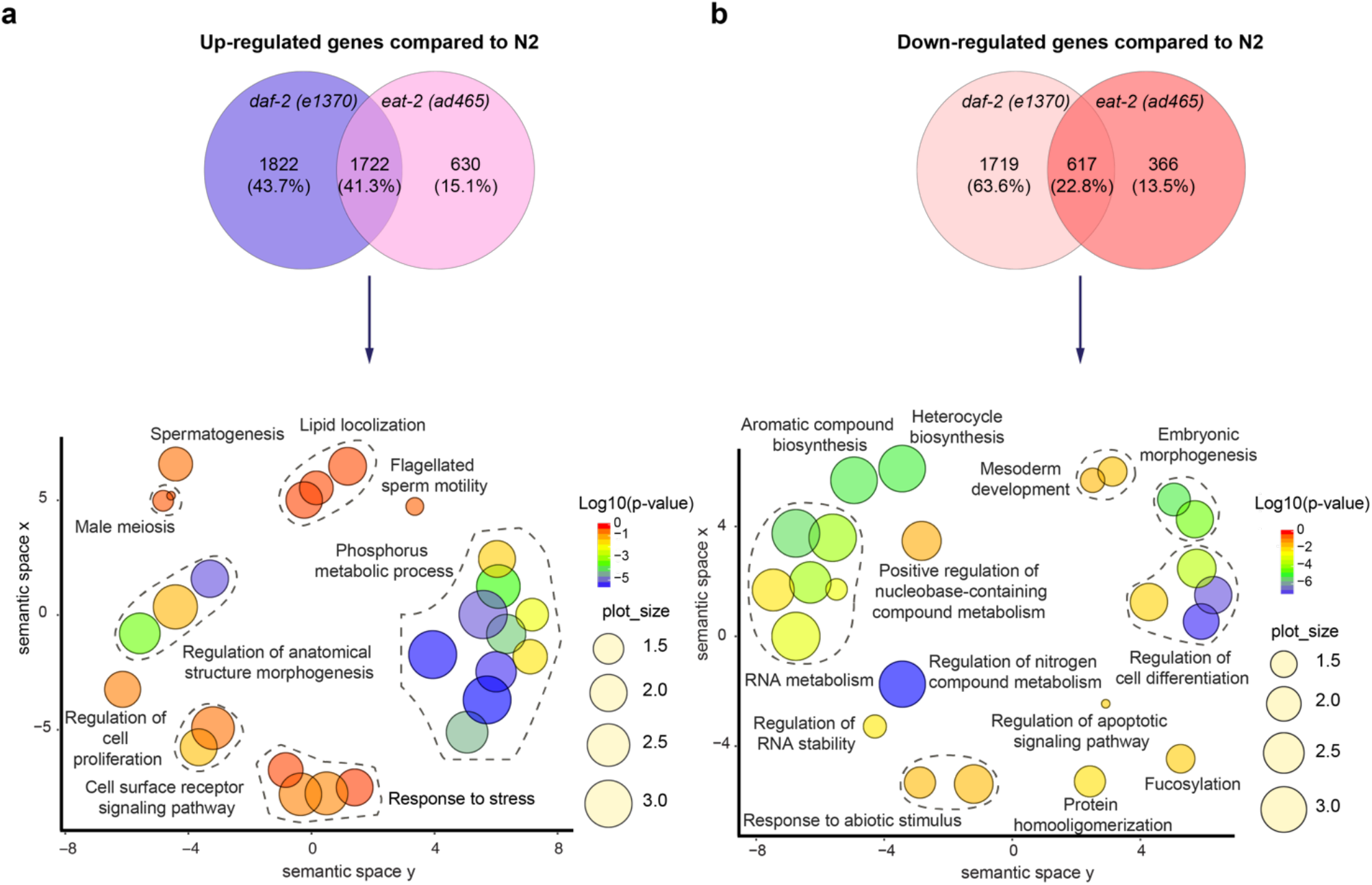
Venn diagram and GO term enrichment analysis comparing *daf-2* and *eat-2* mutants with N2 worms. (**a**) Venn diagram of significantly upregulated genes in both long-lived mutants compared to N2 identified 1722 shared transcripts. GO term enrichment (biological process) of these genes using David and ReviGO highlighted several metabolic pathways such as lipid metabolism and phosphorus metabolic process. (**b**) Venn diagram of significantly downregulated genes in both long-lived mutants compared to N2 identified 617 transcripts. GO term enrichment (biological process) of these genes using David and ReviGO shows an attenuation of process that are involved in biosynthesis of macromolecules such as aromatic compounds, nucleobase-containing compounds and nitrogen compounds. The size of the plot is indicative of the frequency of GO term in the underlying Gene Ontology Annotation(GOA) database (the more the general term, the bigger the plot size); the plots are color-coded according to significance (log_10_-transformed). GO terms belonging to the same cluster were grouped and circled in dark grey dashed line. The GO term at the highest hierarchy level of the cluster is denoted adjacent to the respective dashed line circle.

**Table 1:** KEGG pathway enrichment analysis of the 1722 up-regulated genes and 617 down-regulated genes of both long-lived mutants compared to N2.

| **Upregulated pathways** | **p-value** | **Fold Enrichment** | **Bonferroni** | **Benjamini** | **FDR** |
| --- | --- | --- | --- | --- | --- |
| cel00071:Fatty acid degradation | 7.74E-06 | 13.42 | 4.10E-04 | 4.10E-04 | 7.55E-03 |
| cel01100:Metabolic pathways | 3.97E-05 | 2.47 | 2.10E-03 | 1.05E-03 | 3.88E-02 |
| cel01130:Biosynthesis of antibiotics | 1.00E-04 | 4.87 | 5.30E-03 | 1.77E-03 | 9.78E-02 |
| cel00980:Metabolism of xenobiotics by cytochrome P450 | 4.24E-04 | 13.18 | 2.22E-02 | 5.60E-03 | 4.12E-01 |
| cel00982:Drug metabolism - cytochrome P450 | 6.71E-04 | 11.71 | 3.50E-02 | 7.09E-03 | 6.53E-01 |
| cel01230:Biosynthesis of amino acids | 9.37E-04 | 7.44 | 4.85E-02 | 8.25E-03 | 9.10E-01 |
| cel01212:Fatty acid metabolism | 1.71E-03 | 9.17 | 8.69E-02 | 1.29E-02 | 1.66 |
| cel00350:Tyrosine metabolism | 2.81E-02 | 11.00 | 7.79E-01 | 1.72E-01 | 24.26 |
| cel00260:Glycine, serine and threonine metabolism | 3.53E-02 | 9.73 | 8.51E-01 | 1.91E-01 | 29.57 |
| cel01220:Degradation of aromatic compounds | 4.53E-02 | 42.17 | 9.15E-01 | 2.18E-01 | 36.41 |
| cel00290:Valine, leucine and isoleucine biosynthesis | 4.53E-02 | 42.17 | 9.15E-01 | 2.18E-01 | 36.41 |
| cel00270:Cysteine and methionine metabolism | 4.87E-02 | 8.16 | 9.29E-01 | 2.14E-01 | 38.58 |
| **Downregulated pathways** | **p-value** | **Fold Enrichment** | **Bonferroni** | **Benjamini** | **FDR** |
| cel04350:TGF-beta signaling pathway | 6.10E-06 | 34.34 | 9.15E-05 | 9.15E-05 | 4.24E-03 |
| cel04310:Wnt signaling pathway | 4.95E-05 | 20.49 | 7.42E-04 | 3.71E-04 | 3.44E-02 |
| cel04141:Protein processing in endoplasmic reticulum | 6.60E-05 | 11.30 | 9.89E-04 | 3.30E-04 | 4.59E-02 |
| cel04120:Ubiquitin mediated proteolysis | 2.93E-03 | 12.13 | 4.30E-02 | 1.09E-02 | 2.02 |

### Metabolomics analysis uncovers distinct metabolic changes in both long-lived mutants

In order to provide novel insight into the relationship between longevity and metabolism, we characterized the metabolic fingerprints of the long-lived mutants by performing semitargeted metabolomics with a focus on polar metabolites (Fig. 3). Polar metabolite profiles of the two long-lived mutants and wild-type N2 worms were analyzed using unsupervised PCA analysis which demonstrated marked differences in metabolite profiles between the two long-lived mutants and N2 worms (Fig. 3a). The long-lived mutants differed in a number of metabolic features compared to N2 worms (Fig. 3b). In total, we observed fourteen metabolites that were significantly altered in both long-lived mutants compared to N2 worms. We observed a lowered level of three metabolites in both long-lived mutants when compared to N2 worms, including the glycerolipid intermediate glycerol-3P, the nucleobase adenine, and the purine base xanthine. Additionally, the nucleotide adenosine monophosphate (AMP) was increased in both long-lived mutants. Two metabolites—the TCA intermediate succinate and adenosine diphosphate (ADP)—were significantly changed in both long-lived mutants, although both metabolites were lowered in *daf-2* mutants and increased in *eat-2* mutants. Furthermore, we observed distinct metabolite signatures unique for either of the two long-lived mutants. For instance, we detected three metabolites that are uniquely changed for *daf-2* mutant, including the TCA cycle intermediate citrate/isocitrate, the pentose phosphate pathway (PPP) intermediate gluconate, and glycolytic intermediate phosphoenolpyruvate, which showed a lowered level compared to N2 worms (Supplementary Table S2). In contrast, all three metabolites that were detected specifically for *eat-2* mutants showed an increased level compared to those in N2 worms, including two pyrimidine metabolism intermediates—uridine monophosphate (UMP) and uracil—and one purine metabolism intermediate—guanosine monophosphate (GMP) (Fig. 3b and Supplementary Table S2).

**Figure 3.**
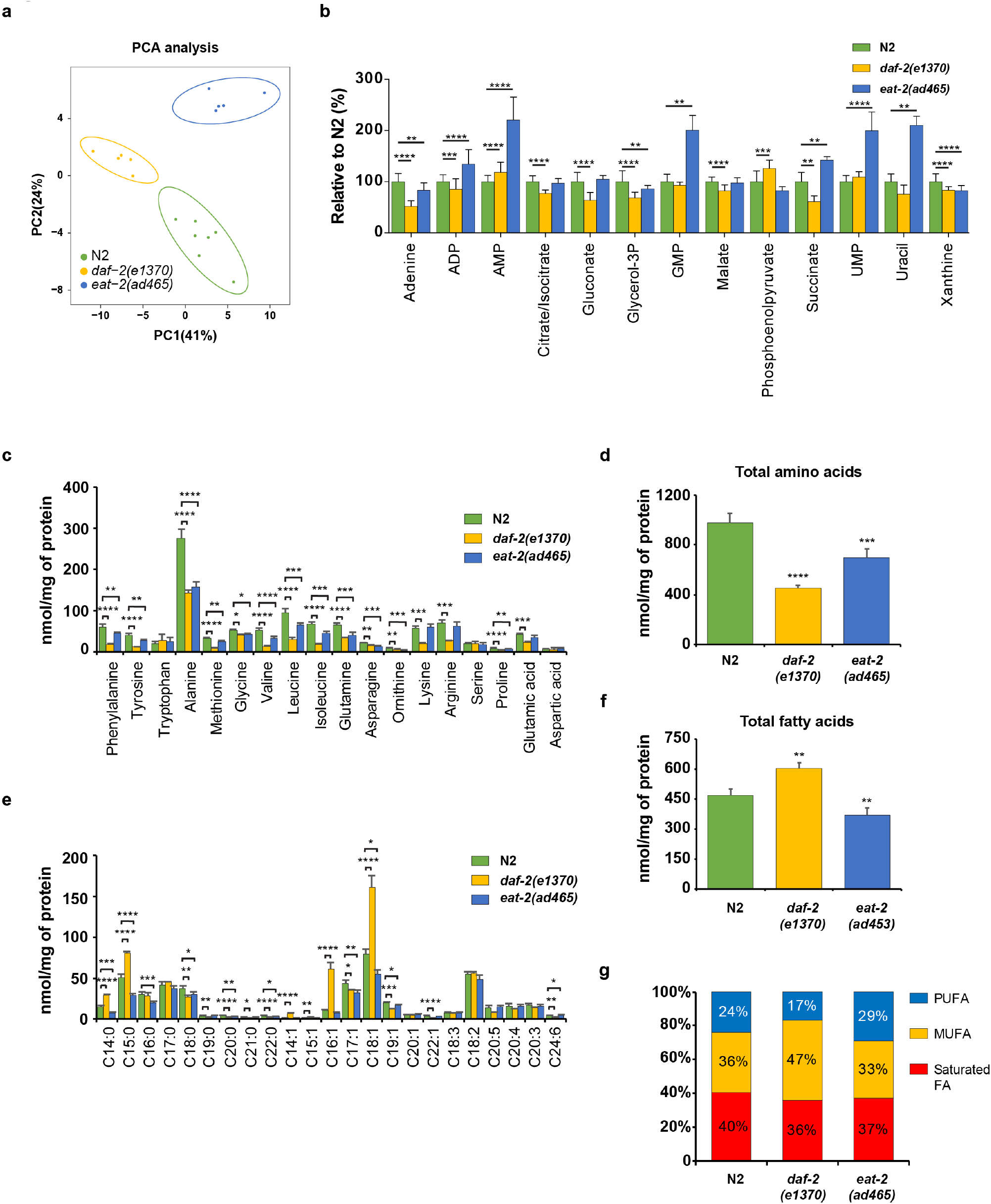
Metabolite signatures of the long-lived mutants *daf-2* and *eat-2*. (**a**) Principle Component Analysis (PCA) analysis score plot showing group separation based on polar metabolite profiles in both long-lived mutants compared to N2 worms. (**b**) Polar metabolite profiles of *daf-2* and *eat-2* mutants compared to N2 worms (FDR<0.05). Several shared metabolic signatures were detected, including increased levels of AMP, decreased levels of glycerol-3P, adenine, and xanthine. Unique metabolite signatures for each long-lived mutant were also observed, for the *daf-2* mutant: increased levels of phosphoenolpyruvate, and decreased levels of citrate/isocitrate, gluconate, ADP, succinate and malate; for the *eat-2* mutant: increased levels of succinate, UMP, uracil and GMP, and decreased ADP. (**c**) Most amino acid species in both long-lived mutants were lowered compared to N2 worms. (**d**) Total amino acids in both *daf-2* and *eat-2* mutants were significantly lower compared to that in N2 worms (**e**) A number of abundant fatty acids were increased in *daf-2* mutants compared to N2 worms, whereas most fatty acids in *eat-2* mutants were decreased compared to N2. (**f**) Total fatty acids in *daf-2* mutants were significantly elevated while those in *eat-2* mutants were significantly decreased when compared to those in N2 worms. (**g**) Mono-unsaturated fatty acids (MUFA) were more abundant in *daf-2* mutants and polyunsaturated fatty acids (PUFAs) were more abundant in *eat-2* mutants compared to N2. The bar graphs depict mean±SD. Significance was calculated using One-way ANOVA; *p ⩽ 0.05; **p ⩽ 0.01; ***p ⩽ 0.001; **** p ⩽ 0.0001.

To expand on the characterization of metabolites in long-lived mutants, we measured amino acids and fatty acids using our quantitative mass spectrometry (MS) platform^18^. We observed a decrease in most individual amino acids in the two long-lived mutants compared to N2 worms (Fig. 3c). The decrease was much more pronounced in *daf-2* mutants, however, and some amino acids did not change in *eat-2* mutants at all (Fig. 3c). The total abundance of amino acids was significantly decreased in both long-lived mutants (Fig. 3d). These results suggest the reorganization of amino acid metabolism in the two long-lived mutants. From the transcriptional data set analysis, we identified several pathways involved in lipid metabolism, including the KEGG pathway terms fatty acid degradation and fatty acid metabolism, and the GO terms lipid localization and lipid storage (Fig. 2, Table 1). We therefore examined the specific lipid signatures in the two long-lived mutants (Fig. 3e). We only observed two changes in fatty acid levels that were shared between both long-lived mutants, namely lowered levels of saturated fatty acid C18:0 and odd-chain monounsaturated fatty acid C17:1. In the *daf-2* mutants only, we observed an elevated level for several saturated fatty acids and monounsaturated fatty acids (MUFAs), including C14:0, C15:0, C14:1, C15:1, C16:1, and C18:1 (Fig. 3e and Supplementary Table S2). Some fatty acids were decreased in *daf-2* mutants, including saturated fatty acids C19:0, C20:0, C21:0, C22:0, MUFA C22:1, and PUFA C24:6. In *eat-2* mutants, fatty acid levels were generally lower, such as the saturated fatty acids C14:0, C15:0, C16:0, C17:0, and the MUFA C18:1. In addition, the total amount of fatty acids in *daf-2* mutants was significantly increased compared to N2 worms, while this was opposite in *eat-2* mutants (Fig. 3f). Interestingly, *daf-2* and *eat-2* mutants showed a distinct lipid saturation; the former has a higher proportion of MUFAs compared to the other two strains, and the latter has more PUFAs (Fig. 3g). Taken together, these results suggest a distinct utilization of lipids in both long-lived mutants and a tight regulation of energy metabolism in these animals.

### Cross-omics analysis highlights important nodes in central carbon metabolism

Following the changes in polar metabolites, we then focused on elucidating key nodes of central carbohydrate metabolism and nucleotide metabolic pathways. We integrated the polar metabolite profiles together with the gene expression profiles for both long-lived mutants and plotted both omics datasets in the known pathways and interactions annotated in KEGG (Fig. 4)^21^. In the glycolysis/gluconeogenesis pathway, we observed an upregulated expression for *pfk-1.2* in both long-lived mutants (Fig. 4). This gene encodes phosphofructokinase, which is one of the major regulatory enzymes of glycolysis that irreversibly catalyze ß-D-fructose-6P to ß-D-fructose-1,6P_2_. We detected downregulation of *gpd-4* (glyceraldehyde-3-phosphate dehydrogenase) in both long-lived mutants, however, this had only a minor influence on the concentrations of glycolytic intermediates in the following steps. In addition, except *pyk-1* (pyruvate kinase) that regulates the final step of glycolysis, expression levels of genes encoding the transitional glycolytic enzymes, including *pgk-1* (phosphoglycerate kinase), *ipgm-1* (phosphoglycerate mutase), and *enol-1* (enolase) remained unaltered in both long-lived mutants. Interestingly, we detected an elevated level of phosphoenolpyruvate in *daf-2* mutants whereas a decreased level of phosphoenolpyruvate is observed in *eat-2* mutants compared to N2 worms. This may be explained by the discovery that the expression level of *pck-2* (phosphoenolpyruvate carboxykinase) was upregulated in *daf-2* mutants while down-regulated in *eat-2* mutants. Furthermore, we observed an increased expression of *pyk-1* (pyruvate kinase), which may explain the elevated levels of pyruvate in *daf-2* mutants while it remained unchanged in *eat-2* mutants. Another gene that encodes a key glycolytic enzyme is hexokinase, *hxk-1*, which was downregulated in *daf-2* mutants, possibly explaining our observations that ß-D-fructose-1,6P_2_ was lowered in *daf-2* mutants whereas in *eat-2* mutants *hxk-1* expression was elevated compared to N2 worms.

**Figure 4.**
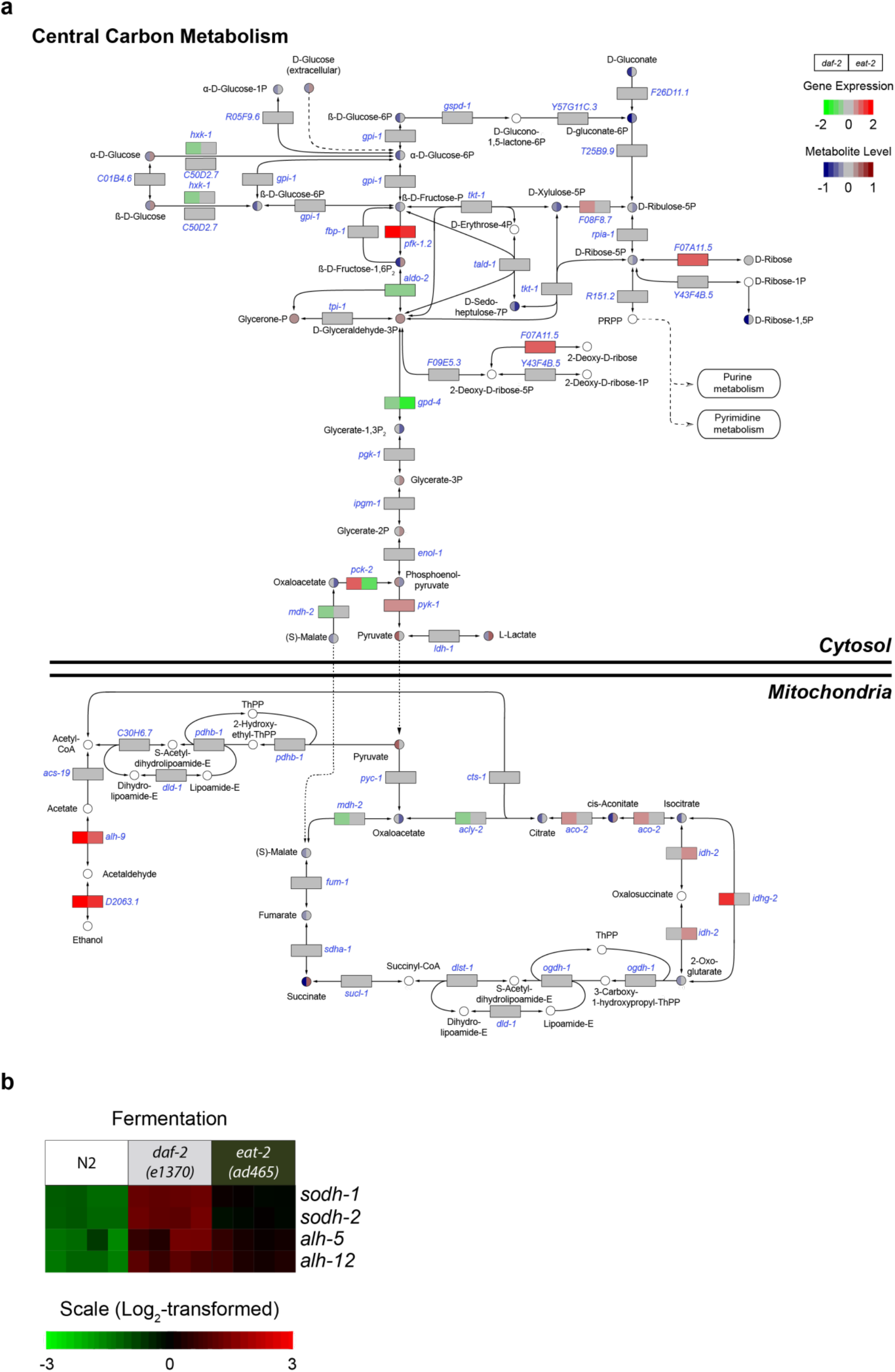
Central carbon metabolism pathways in *daf-2* and *eat-2* mutants. (**a**) Data from the gene expression profile (rectangles) and metabolite profile (circles) were color-coded and integrated in three major pathways of central carbon metabolism, including glycolysis/gluconeogenesis, the pentose phosphate pathway and TCA cycle. The majority of genes involved in glycolysis/gluconeogenesis remained unaltered in both long-lived mutants. Glycolysis/gluconeogenesis intermediates remained low in both long-lived mutants, except the final product of this pathway, i.e. pyruvate, which accumulated in *daf-2* mutants. Genes coding for PPP enzymes were mostly unchanged and PPP intermediates were lowered in both long-lived mutants. Alcohol fermenting genes also showed increased expression in both long-lived mutants. The majority of genes involved in the TCA cycle were unchanged and TCA intermediates were significantly lowered in *daf-2* mutants. TCA intermediates were unchanged in *eat-2* mutants except for elevated cis-aconitate and succinate levels. White circles are metabolites that were not measured. (**b**) Gene expression profiles of genes that are involved in fermentative metabolism. Each column represents one biological replicate.

The PPP is also involved in cytosolic central carbon metabolism. Transcript levels of most PPP genes in both long-lived mutants showed no significant differences compared to those in N2 worms, including *gdpd-1* (glucose-6-phosphate dehydrogenase), *Y57G11C.3* (6-phosphogluconolactonase), *T25B9.9* (phosphogluconate dehydrogenase), *tkt-1* (transketolase), *rpia-1* (ribose 5-phosphate isomerase A), *tald-1* (transaldolase 1), *R151.2* (phosphoribosylpyrophosphate synthetase 1), and *Y43F4B.5* (phosphoglucomutase 2) (Fig. 4). We did, however, detect a significant upregulation in the expression of *F07A11.5*, an orthologue of human ribokinase (RBKS), in both long-lived mutants. Moreover, we observed an upregulation of *F08F8.7*, a gene encoding an enzyme that is predicted to have ribulose-phosphate 3-epimerase activity, in *daf-2* mutants compared to N2. Most PPP metabolite concentrations in both long-lived mutants remained either unchanged or relatively lower compared to N2. Taken together, the selective increased transcription levels of PPP genes and low levels of PPP intermediates suggests a lowered flux in this pathway in both long-lived mutants compared to N2 worms.

In both long-lived mutants, we observed a strong upregulation of genes encoding enzymes involved in alcohol fermentation, including *alh-5* (aldehyde dehydrogenase 3 family member B2), *alh-12* (aldehyde dehydrogenase 9 family member A1), *sodh-1* (sorbitol dehydrogenase family), and *sodh-2* (sorbitol dehydrogenase family) (Fig. 4A and 4B, Supplemental Table S1). The substrates of these dehydrogenases remain unclear, although SODH-1 has been reported to catalyze the breakdown of ethanol ^22^. In addition, SODH-1 was predicted to have oxidoreductase function and its expression can be regulated by several essential metabolic genes, including *daf-16* (FOXO), *lbp-5* (lipid binding protein), *mdt-15* (mediator subunit), and *pept-1* (peptide transporter)^23-25^. Hence, the known biological activity of SODH-1 may include more than its function in fermentative metabolism alone. Furthermore, upregulation of *alh-5* and *alh-12* may lead to higher intracellular levels of acetate which, together with the observed unchanged expression of *pdhb-1* (pyruvate dehydrogenase beta), may partly explain the low abundance of citrate that we observed in *daf-2* mutants and unaltered levels in *eat-2* mutants, respectively.

As mentioned above, transcript profiles of both long-lived mutants did not show pronounced changes in mRNA levels for TCA cycle enzymes, and this pathway was therefore not identified in the KEGG pathway and GO term enrichment analyses (Table 1, Fig. 4). However, a few genes that encode key enzymes in the TCA cycle were differentially regulated in *daf-2* and *eat-2* mutants compared to N2. Specifically, *aco-2* (aconitase) and *idhg-2* (isocitrate dehydrogenase gamma), were upregulated, and *mdh-2* (malate dehydrogenase) and *acly-2* (ATP citrate lyase) were downregulated in *daf-2* mutants, and *idh-2* (isocitrate dehydrogenase) was upregulated in *eat-2* mutants (Fig. 4). Furthermore, we also observed low levels of TCA cycle intermediates in both long-lived mutants, except for cis-aconitate and succinate that were relatively high in the *eat-2* mutant compared to N2. These findings suggest that energy metabolism in both long-lived mutants is differentially regulated compared to that in N2 worms. Moreover, the TCA cycle may be particularly impaired in *daf-2* mutants, as shown by the high level of pyruvate and reduced levels of TCA intermediate metabolites.

### Pyrimidine metabolism is downregulated in both long-lived mutants

In characterizing the metabolic fingerprints of both long-lived mutants compared to N2 worms, we observed marked changes in nucleotides that are involved in the pyrimidine metabolic pathway. This is in line with previous work showing that the regulation of pyrimidine metabolism is *daf-16*-dependent^14^. Indeed, we observed a general decrease of pyrimidine metabolic gene transcripts in both long-lived mutants, especially in *daf-2* mutants (Fig. 5). For instance, *rpb-6*, a gene encoding a human orthologue of the RNA polymerase II subunit F, was significantly downregulated in both long-lived mutants and likely resulted in the low levels we observed for UTP and CTP (Fig. 5). We also detected low expression of *ndk-1* (nucleoside diphosphate kinase), which may explain the reduced levels of dTDP in the *eat-2* mutants. Another downregulated pyrimidine metabolic gene was *rnr-1*, which encodes the large subunit of ribonucleotide reductase and catalyzes the irreversible conversion of CDP and UDP to dCDP and dUDP, respectively.

**Figure 5.**
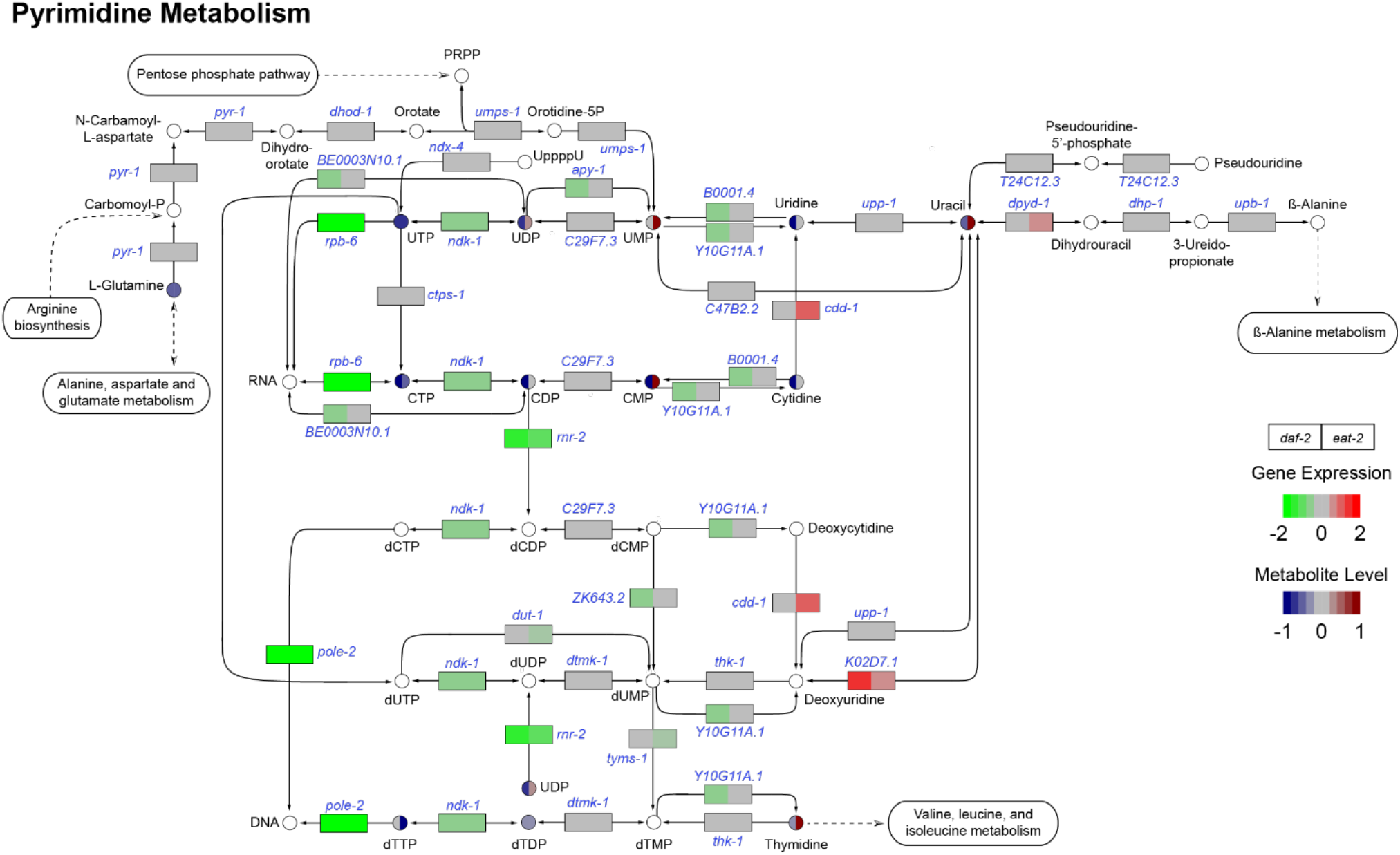
Pyrimidine metabolism pathways in *daf-2* and *eat-2* mutants. Genes involved in the pyrimidine metabolism in both long-lived mutants were significantly down-regulated compared to those in N2 worms. Only a few genes involved in pyrimidine degradation were upregulated. In line with the expression, most pyrimidine intermediates were low abundant in both long-lived mutants. White circles are metabolites that were not measured.

Overall, we observed decreased concentrations of pyrimidine intermediates in *daf-2* mutants that corroborates a down-regulation of the majority of genes involved in pyrimidine metabolism. In contrast, although many pyrimidine metabolic genes have lowered expressions in the *eat-2* mutants, we observed elevated concentrations of the pyrimidine intermediate metabolites compared to N2, including UDP, UMP, CMP, uracil, and thymidine (Fig. 5). Elevated levels of uracil may be associated with increased expression levels of *dpyd-1* (dihydro-pyrimidine dehydrogenase) and *K02D7.1* (purine nucleoside phosphorylase). Overall, our results from the cross-omics analysis revealed a down-regulation of pyrimidine metabolic pathways in both long-lived mutants compared to N2 worms.

### Purine de novo synthesis pathways were enhanced in both long-lived mutants

We next explored the purine metabolic pathways, including *de novo* synthesis, the salvage pathways and purine degradation (Fig. 6). Most genes involved in *de novo* purine biosynthesis pathways, including *ppat-1* (phosphoribosyl pyrophosphate amidotransferase), *F38B6.4* (phosphoribosylamine-glycine ligase and phosphoribosyl-formylglycinamidine cyclo-ligase), *pacs-1* (phosphoribosylamino-imidazole carboxylase and phosphor-ribosylaminoimidazole-succinocarboxamide synthase), and *atic-1* (5-aminoimidazole-4-carboxamide ribonucleotide formyltransferase/IMP cyclo-hydrolase), were upregulated in both long-lived mutants. We observed a low level of ribose-5P in *eat-2* mutants compared to N2 and *daf-2* animals. Interestingly, the abundance of 5-formamidoimidazole-4-carboxamide-ribotide (FAICAR) and inosine monophosphate (IMP), the intermediates at the final two steps of purine ring synthesis, were decreased in *daf-2* mutants but elevated in *eat-2* mutants compared to N2 worms. Genes encoding enzymes involved in the salvage pathway are shared between purine and pyrimidine metabolism and we therefore observed a similar gene expression profile: most genes were the either downregulated or remain unchanged in both long-lived mutants. Similar to IMP, the levels of other purine nucleotides, including GDP, GMP, AMP, and ADP were elevated in *eat-2* mutants, while decreased in *daf-2* mutant. GTP and ATP remained at low levels in both long-lived mutants. Notably, genes involved in purine degradation were upregulated in both long-lived mutants. Most nucleotide metabolic intermediates displayed complex patterns: some showed comparable concentrations to their upstream substrates, such as xanthine and hypoxanthine; guanine showed an opposite level compared to guanosine and GMP, which was elevated in *daf-2* mutants and lowered in *eat-2* mutants compared to N2 worms (Fig. 6). Overall, this result suggests an up-regulation of purine metabolism, including de novo synthesis, salvage pathways and degradation.

**Figure 6.**
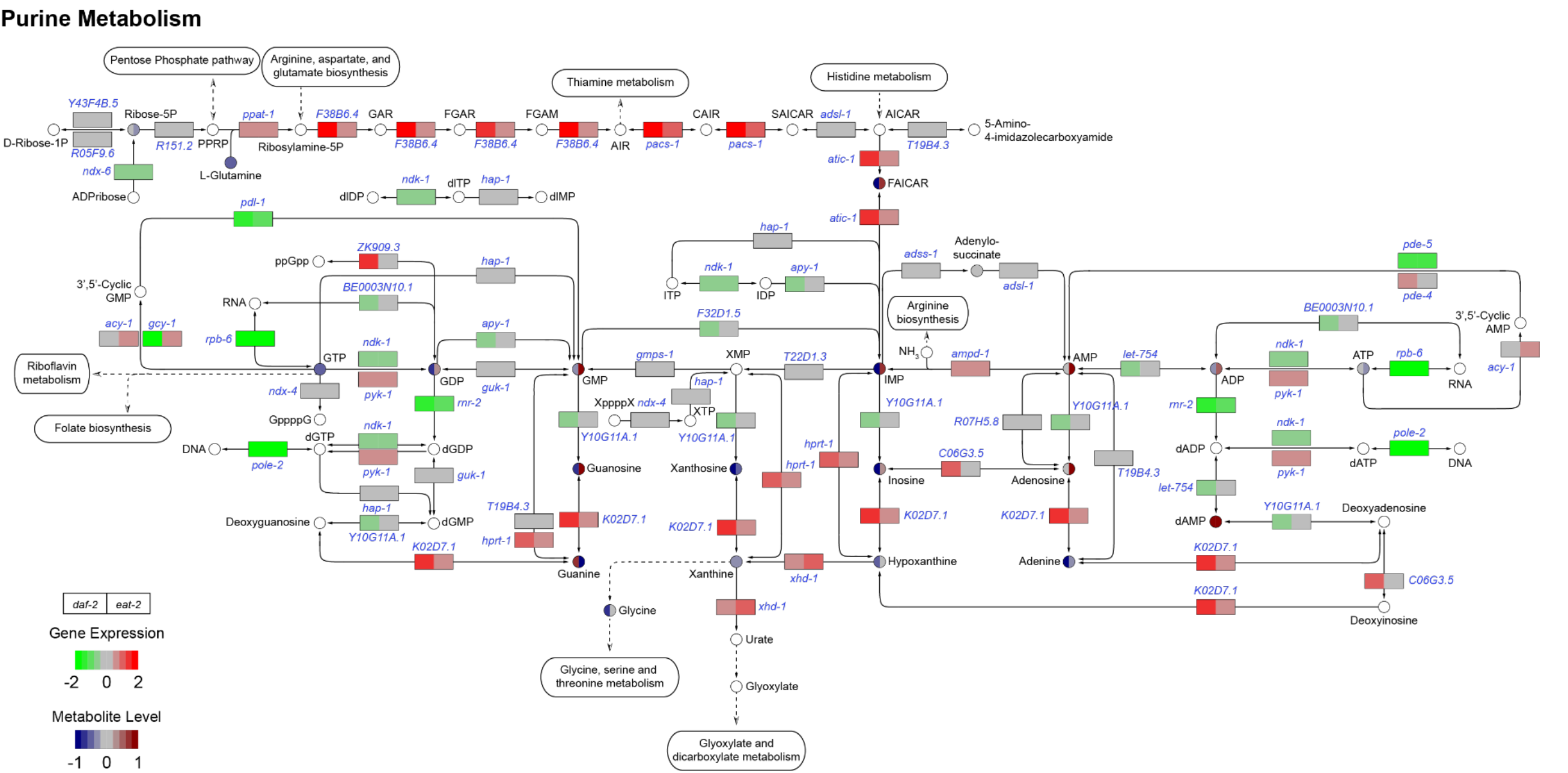
Purine metabolism pathways of *daf-2* and *eat-2* mutants. The majority of genes involved in the three major pathways (de novo synthesis, salvage, and degradation) of purine metabolism were significantly upregulated in both *daf-2* and *eat-2*. Three metabolites showed similar changes in both mutants, i.e. AMP, adenine, and xanthine. Most purine metabolism intermediates were decreased in *daf-2* mutants, except guanine; in *eat-2* mutants the differences in purine intermediate abundance is more diverse. White circles are metabolites that were not measured.

### Other altered metabolic pathways related to amino acids changes in the long-lived mutants

In addition, we also analyzed cross-omics data for the altered amino acid profiles in both long-lived mutants. Because we observed lower levels of methionine and glycine, which are involved in one carbon metabolism, we checked the changes at the transcriptional level. Indeed, genes involved in one carbon metabolism, such as *atic-1, dao-3, F38B6.4, gcst-1*, and *K07E3.4*, were significantly upregulated in *daf-2* mutants and *alh-3, atic-1, F38B6.4, gcst-1*, and *K07E3.4 were* upregulated in *eat-2* mutants (Fig. 7a and Supplementary Table S1). This suggests that one carbon metabolism may be enhanced in both long-lived mutants. To gain insight into BCAA metabolism modulation, we evaluated the transcriptomic data sets and found that the majority of genes involved in the BCAA degradation pathway were altered in their expression levels in both long-lived mutants. In *daf-2* mutants, *Y44A6D.5, acdh-8, alh-12, F54C8.1, gta-1, hacd-1*, and *mce-1* were upregulated whilst in *eat-2* mutants, approximately 50% of the DEGs were upregulated (e.g. *bcat-1)* and the rest were downregulated, compared to N2 worms (Fig. 7b and Supplementary Table S1). The distinct alterations at the transcriptional level may explain why the level of BCAAs in *eat-2* mutants were lowered compared to N2, and especially drastically reduced in *daf-2* mutants. From the functional annotation clustering for KEGG pathway terms, we observed a significant enrichment for tyrosine metabolism, glycine metabolism and degradation of aromatic compounds. Together with the previous observed lowered level of these amino acids in both *daf-2* and *eat-2* mutants, these data suggest a shift from carbon to amino acid catabolism as alternative energy source in both long-lived mutants.

**Figure 7.**
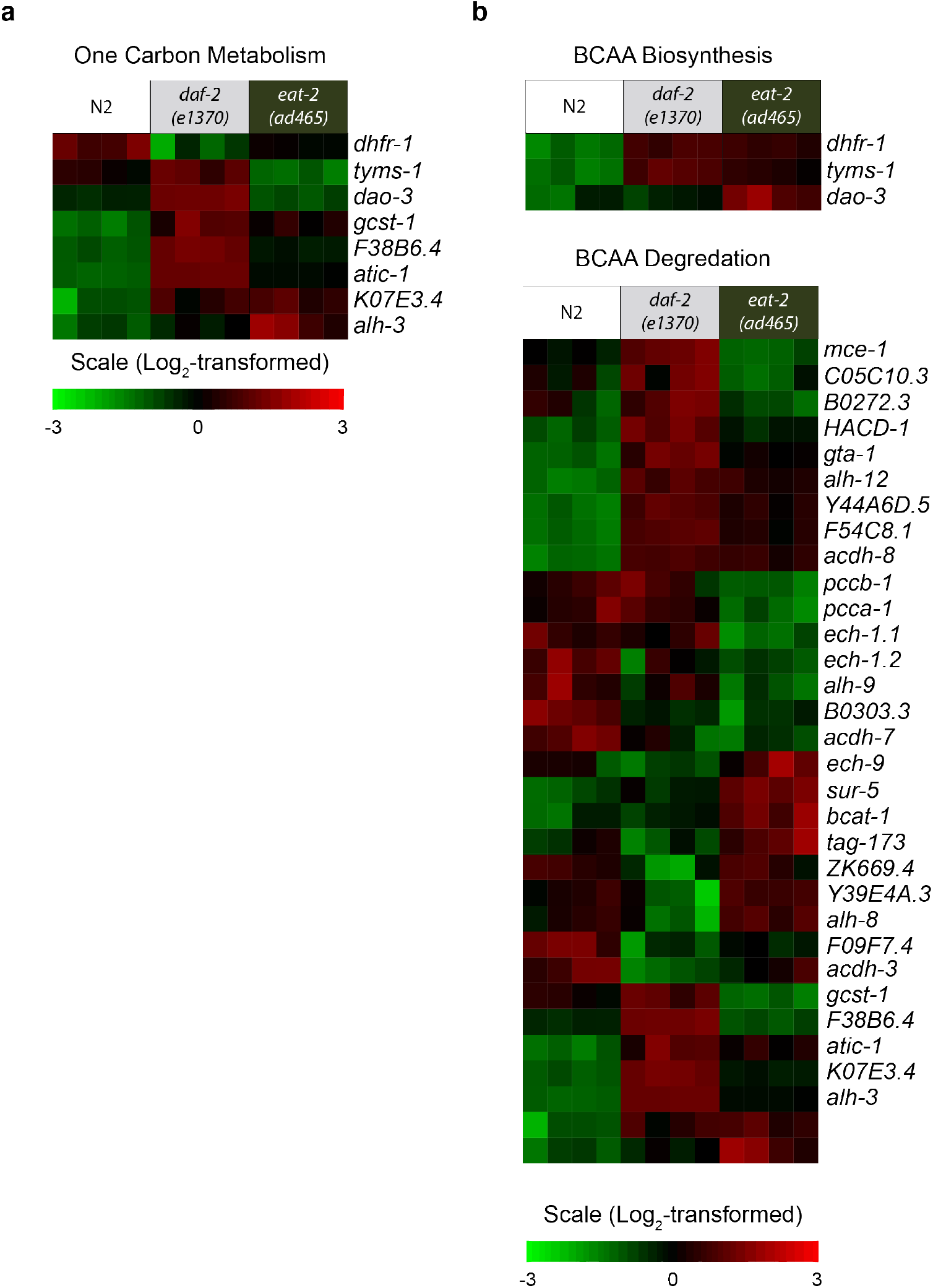
Heat maps showing transcriptional changes in specific gene sets. Gene expression profiles of gene sets that are involved in one carbon metabolism (**a**), and branched-chain amino acid (BCAA) metabolism (**b**). Each column represents one biological replicate.

## Discussion

Impaired IIS and caloric restriction are two well-known longevity interventions that operate in parallel or through partially independent mechanisms^3^. Previous studies with whole-genome transcript profiling of long-lived mutants have discovered differential expression of gene sets that are involved in lipid metabolism, oxidative stress response and heat shock stress response^23,26,27^. Because most of these genes play important roles in energy metabolism, recent studies have put great effort into characterization of the metabolic consequences in these long-lived mutants^14,28,29^. However, a correlation between the enzyme regulation at a transcriptional level and their metabolite concentrations remain unclear, especially with regard to energy metabolism^29^. Although a number of essential regulators have been identified at the transcriptomic and metabolomic level of both long-lived mutants, the common mechanisms that underlie metabolic control of longevity in *daf-2* and *eat-2* mutants have not been elucidated. One possible reason is a lack of comprehensive metabolomics analysis and multi-layer omics platforms. In this study, we hence present an in-depth identification of changes at both the transcript and metabolite level to acquire comprehensive information of potentially common metabolic longevity pathways shared in two long-lived mutants. By analyzing the gene expression profile of *daf-2* and *eat-2* mutants, we have identified a number of genes in both animals that similarly regulate regions of the metabolic network. The analysis of KEGG pathways and GO terms on genes upregulated in both long-lived mutants showed a significant enrichment of genes involved in regulation of lipid metabolism (lipid storage and lipid localization) and amino acid metabolism (tyrosine, glycine, serine threonine, methionine and BCAAs). Conversely, biosynthesis of macromolecules, such as aromatic compounds and nucleobase-containing compounds, was downregulated in both long-lived mutants. Most of the transcriptomics findings are in a good agreement with previous studies in *eat-2*^27^ or *daf-2*^30^. Collectively, these findings suggest a metabolic shift in both mutants where amino acids may be used as a primary energy source. In order to determine the metabolic consequence of these altered pathways and identify novel metabolite signatures for longevity, we also collected metabolite profiles from both *daf-2* and *eat-2* mutants using semi-targeted and targeted metabolomics platforms. We observed several novel metabolite features shared in *daf-2* and *eat-2* mutants, including elevated levels of the purine salvage pathway intermediate (AMP) and reduced levels of the glycerolipid intermediate (glycerol-3P), and purine degradation intermediates (adenine and xanthine). These findings suggest the existence of shared metabolite profiles and common alterations in biological processes that are required for longevity in both long-lived mutants. In addition, we also identified novel metabolite features specifically present in either of these long-lived mutants. Several central carbon metabolism intermediates were lowered in the *daf-2* mutants, including succinate, citrate/isocitrate, gluconate and phosphoenolpyruvate, suggesting an altered central carbon metabolism in *daf-2* animals. Specific metabolic signatures for the *eat-2* mutant were elevated levels of succinate and both pyrimidine and purine metabolism intermediates, suggesting an altered regulation in nucleotide metabolism. From the targeted metabolomics analysis, we observed an overall low amino acid abundance in both long-lived mutants, which was consistent with our gene expression data and comparable with previous studies^31,32^. Several of the amino acids are used as an alternative supply for TCA cycle intermediates, including alanine, glutamine, tyrosine, and phenylalanine, and were lowered in both long-lived mutants. Together with the enriched KEGG pathways, these findings suggest a metabolic shift in both long-lived mutants and a strong involvement of amino acid catabolism as an alternative carbon source for the TCA cycle.

In the fatty acid profiles of these two long-lived mutants we observed two fatty acids that showed a similar profile, i.e. lowered concentration of C18:0 and C17:1. In *daf-2* mutants, we detected relatively higher MUFAs levels—especially due to high levels of C16:1 and C18:1— while in *eat-2* mutants, we observed higher PUFAs levels. Because we identified a significant enrichment of the lipid storage pathway in upregulated genes in both long-lived mutants, we further analyzed the gene clusters involved in the lipid metabolism. We observed an upregulation of genes involved in fatty acid biosynthesis and a downregulation of those involved in fatty acid degradation in both long-lived mutants.

To further uncover potentially shared metabolic pathways that are important for longevity in both *daf-2* and *eat-2* mutants, we integrated transcriptome and metabolome datasets based on known KEGG pathways involved in central carbon metabolism, pyrimidine and purine metabolism. We analyzed three major pathways involved in central carbon metabolism, including glycolysis/gluconeogenesis, the pentose phosphate pathway (PPP), and TCA cycle. We detected a distinct regulation at both transcript and metabolite levels involved in glycolysis/gluconeogenesis in *daf-2* and *eat-2* mutants. Proteomics analysis previously identified a set of proteins involved in amino acid metabolism, carbohydrate metabolisms and ROS detoxification that was significantly altered in *daf-2* mutants, although such proteins do not causally contribute to lifespan extension of *daf-2* mutants^33^. In *eat-2* mutants, genes encoding enzymes that are involved in glycolysis and gluconeogenesis were downregulated, which is consistent with proteomics analyses^34^. Because we have observed lower amino acids levels and increased fatty acid levels in our metabolomics analysis, it is likely that amino acids serve as an alternative energy source for *daf-2* mutants^29,35,36^. Similarly, a previous proteomics study on *eat-2* mutants showed a subtle shift from carbohydrate to fatty acid metabolism^34^, which is also partly reflected in our data. Additionally, our data pinpoint amino acids as an alternative nutrient source in *eat-2* mutants. Overall, our data suggest that both long-lived mutants display a large degree of metabolic flexibility, which allows them to cope better with the available energy source and maintain metabolic homeostasis.

Our results revealed a downregulation of pyrimidine anabolic pathways and an upregulation of pyrimidine degradation in both long-lived mutants. Although regulation of pyrimidine anabolism at gene expression level was similar in both long-lived mutants, we especially detected high levels of several pyrimidine intermediates in *eat-2* mutants. In contrast, purine metabolism, including *de novo* synthesis, salvage and degradation pathways, was upregulated in both long-lived mutants at the transcriptional level. At the metabolite level, although *daf-2* and *eat-2* mutants showed distinct features, two intermediates of purine degradation were lowered in both long-lived mutants (adenine and xanthine), suggesting a novel metabolic signature shared between the two long-lived mutants. Interestingly, downregulation of pyrimidine metabolism was previously reported in the long-lived *glp-1* mutant in a *daf-16* dependent manner, suggesting the downregulated pyrimidine metabolism could be an important downstream target in both *daf-2* and *eat-2* mutants^14^.

In conclusion, the integration of transcriptomics and metabolomics analyses provides an extensive dataset of changes resulting from impaired IIS and CR in *C. elegans*. Importantly, although *daf-2* and *eat-2* mutants have extended lifespan through disparate mechanisms, impaired IIS and CR share transcriptome and metabolome signatures that likely facilitate longevity. These novel findings provide important insights into the relationship between metabolism and longevity and help pave the way towards metabolic and pharmacological interventions that may enhance longevity.

## Materials and methods

### C. elegans strains and bacterial feeding strains

The *C. elegans* N2 (Bristol), *daf-2(e1370)* and *eat-2(ad465)*, and *E. coli* OP50 and HT115 strains were obtained from the *Caenorhabiditis* Genetics Center (CGC). Both mutant strains were backcrossed three times and sequenced before use.

### Worm growth conditions and worm pellet collection

*C. elegans* were routinely grown and maintained on nematode growth media (NGM) at 20°C. Worms of each strain were cultured on plates seeded with *E. coli* OP50, then eggs were obtained by alkaline hypochlorite treatment of gravid adults and placed onto plates seeded with *E. coli* HT115. At the young adult stage, each strain was collected by washing them off the culture plates with M9 buffer and the worm pellet was washed with dH_2_O for three times before collection in a 2 mL Eppendorf tube, snap-frozen in liquid N_2_ and lyophilized overnight. For RNA extraction, 500 worms/tube was collected in four replicates per worm strain; for polar metabolite extraction, 2000 worms/tube was collected in five to six replicates; for fatty acid and amino acid extraction, 2000 worms/tube was collected in four replicates.

### Microarray

Total RNA was extracted using the standard Trizol method, followed by phenol-chloroform purification and further processing by OakLabs Gmbh (Germany). In short, RNA quality and quantity was assessed after DNase clean-up using a 2100 Bioanalyzer (Agilent Technologies). All samples had a RNA integrity value of ≥7.2. RNA was amplified and labeled using a Low Input QuickAmp Labeling Kit (Agilent Technologies) and hybridized using the Agilent Gene Expression Hybridization Kit (Agilent Technologies). An ArrayXS-068300 with WormBase WS241 genome build (OakLabs) was used and fluorescence signals were detected by the SureScan microarray Scanner (Agilent Technologies). Data of all samples were quantile normalized using the ranked median quantiles as described previously ^37^. The most significant coding-gene isoform was selected to represent expression for mentioned genes. Fold-change of gene expressions in the mutant strains was characterized as log_2_ (expression_*daf-2 or eat-2 mutant*_/expression_N2_), and the significance (p-value) was calculated using a student’s t-test. The expression of a certain gene in each long-lived mutant with a p-value less than 0.05 and a fold-change either less than -1 or higher than 1 was considered as a significant compared to N2 and selected for further analysis. The overlapping up- or down- regulated genes in the long-lived mutants were selected using Venny diagram 2.1.0^38^. KEGG pathway analysis was performed with David^19^. GO term enrichment analysis was performed with David and ReviGo^20^. The result from ReviGO was downloaded as an R script and images were imported into Illustrator (Adobe) for further editing.

### Metabolomics analysis: polar metabolite measurements

500 μL MQ water, 500 μL MeOH and 1 mL chloroform was added to the dry worm pellet. Thereafter, the worm suspension was homogenized using a TissueLyser II (Qiagen) for 5 min at a frequency of 30 times/sec. After centrifugation at 4°C, ~800 μL of the “polar” top layer was transferred to a new 1.5 mL Eppendorf tube and dried in a vacuum concentrator at 60°C (GeneVac). Dried residues were dissolved in 100 μL MeOH:MQ water (6:4, v/v). For polar metabolite measurements, we used a Thermo Scientific ultra-high pressure liquid chromatography system (Waltman, MA, USA) coupled to a Thermo Q Exactive (Plus) Orbitrap mass spectrometer (Waltman, MA, USA). The auto-sampler was kept at 10°C during the analysis and 5 μL sample was injected on the analytical column. The chromatographic separation was established using a Sequant ZIC-cHILIC column (PEEK 100 x 2.1 mm, 3.0 μm particle size, Merck, Darmstadt, Germany) and held at 15 °C. The flow rate was kept at 0.250 mL/min. The mobile phase was composed of (i) acetonitrile:water (9:1, v/v) with 5 mM ammonium acetate; pH=6.8 and (ii) acetonitrile:water (1:9, v/v) with 5 mM ammonium acetate; pH=6.8, respectively. The LC gradient program was: start with 100% (i) hold 0-3 min; ramping 3-20 min to 36% (i); ramping from 20-24 min to 20% (i); hold from 24-27 min at 20% (i); ramping from 27-28 min to 100% (i); and re-equilibration from 28-35 min with 100% (i). The MS data were obtained in negative ionization mode at full scan range at a mass resolution of 140,000. Interpretation of the data was performed in the Xcalibur software (Thermo scientific, Waltman, MA, USA). α-, β-, D-, & L- metabolites were not distinguished. Statistical analysis was performed using the statistical programming software R combined with packages MixOmics, ropls and ggplot2. Metabolites in either *daf-2* or *eat-2* mutants were determined statistically different from N2 worms using a multi t-test and FDR method of Benjamini and Hochberg using GraphPad Prism, with Q=5%.

### Metabolomics analysis: Fatty acid and amino acid extraction and analysis

Fatty acids were extracted and analyzed as described^18^. In brief, a dry worm pellet was resuspended in 250 μL ice-cold 0.9% NaCl solution. Worm lysate was obtained by adding a 5 mm steel bead to the worm suspension and mixed using a TissueLyser II (Qiagen) for 2x2.5 min at frequency of 30 times/sec, followed by tip sonication (energy level: 40 joule; output: 8 watts) for two times on ice-cold water. Protein quantification was performed with a BCA assay. Worm lysate (containing 150 μg protein) was transferred in a 4 mL fatty acid-free glass vial, and 1 mL of freshly prepared 100% acetonitrile (ACN) / 37% hydrochloric acid (HCl) (4:1, v/v) was added to the lysate, together with deuterium-labeled internal standards. Samples were hydrolyzed by incubating at 90°C for 2 h. After the vials cooled down to room temperature, fatty acids were separated in a hexane layer and then evaporated at 30°C under a stream of nitrogen. Fatty acid residues were dissolved in 150 μL chloroform-methanol-water (50:45:5, v/v/v) solution containing 0.0025% aqueous ammonia, and then transferred to a Gilson vial for ESI-MS analysis.

Amino acids (AAs) were extracted by transferring worm lysate (contains 50 μg of protein) to a 2 mL Eppendorf tube, and 1 mL of 80% ACN plus 20 μL of internal standard mixture was added to the lysate and homogenized by vortexing^18^. Samples were centrifuged and the supernatant was transferred to a 4 mL glass vial and evaporated under a stream of nitrogen at 40°C. After evaporation, AA residue was dissolved in 220 μl of 0.01% heptafluorobutyric acid (v/v in MQ water). Then the suspension was transferred to a Gilson vial for HPLC-MS/MS analysis.

### Cross-omics data analysis

Microarray transcriptomics data and polar metabolite data were integrated using Pathview in R. Metabolic enzymes nodes compromised of multiple enzymes were colored using the sum of all related gene expressions. The first gene listed in KEGG for a certain enzyme was selected for use in the figure.

### Gene expression profile visualization

Heat maps of gene expression profile were plotted using “R2” (Genomics analysis and Visualization platform (<u>http://r2.amc.nl</u>).

## Acknowledgements

We thank the Caenorhabditis Genetic Center, which is funded by NIH Office of Research Infrastructure Programs (P40 OD010440), for supplying *C. elegans* strains. AWG is supported by an AMC PhD Scholarship. Work in the Houtkooper group is financially supported by an ERC Starting grant (no. 638290), and a VIDI grant from ZonMw (no. 91715305).

## Author contributions

AWG, RLS, and RHH conceived and designed the project. AWG, RLS, RK and MvW performed experiments and interpreted data. AWG, RLS and MvW performed bioinformatics. AWG, RLS and RHH wrote the manuscript, with contributions from all other authors.

## Additional Information

### Supplementary information

Supplementary information includes two supplementary table containing the full transcriptomics datasets and metabolomics datasets.

### Data availability

The microarray data is available at Gene Expression Omnibus (accession number: GSE106672)

